# Analysis of single nucleotide variants in CRISPR-Cas9 edited zebrafish embryos shows no evidence of off-target inflation

**DOI:** 10.1101/568642

**Authors:** Marie R Mooney, Erica E Davis, Nicholas Katsanis

## Abstract

Therapeutic applications of CRISPR-Cas9 gene editing have spurred innovation in Cas9 enzyme engineering and single guide RNA (sgRNA) design algorithms to minimize potential off-target events. While recent work in rodents outlines favorable conditions for specific editing and uses a trio design to control for the contribution of natural genome variation, the potential for CRISPR-Cas9 to induce *de novo* mutations *in vivo* remains a topic of interest. In zebrafish, we performed whole exome sequencing (WES) on two generations of offspring derived from the same founding pair: 54 exomes from control and CRISPR-Cas9 edited embryos in the first generation (F0), and 16 exomes from the progeny of inbred F0 pairs in the second generation (F1). We did not observe an increase in the number of transmissible variants in edited individuals in F1, nor in F0 edited mosaic individuals, arguing that *in vivo* editing does not precipitate an inflation of deleterious point mutations.

## Introduction

CRISPR-Cas9 gene editing technology has offered powerful investigative tools and opened new potential avenues for the treatment of genetic disorders. Nonetheless, like preceding technologies, the clinical implementation of CRISPR-Cas9 editing faces potential barriers. These include restricted control over the delivery and activity of the system; immune responses to the system components; and permanent alteration of unintended genomic targets (Ho et al., 2018). In cell culture systems, the alteration of off-target regions decreases precipitously with the use of stringently designed sgRNA sequences and Cas9 enzymes engineered for high specificity (Doench et al., 2016; Fu et al., 2013; Hu et al., 2018), though recent work demonstrates that precise control over the nature of editing even at on-target sites remains challenging (Kosicki et al., 2018). In rodents, these same factors influence the efficiency and specificity of CRISPR-Cas9 editing (Anderson et al., 2018). However, examination of atypical CRISPR-Cas9 influence on organisms remains limited; it is often focused primarily on predicted off-target assessment and is not always agnostic (Varshney et al., 2015).

Here, we evaluated the incidence and transmission of off-target effects in a cohort of CRISPR-Cas9 edited zebrafish embryos derived from the same founding pair. Using 52 zebrafish embryos from the same clutch targeted with sgRNAs with variable on-target efficiency, we whole-exome sequenced DNA from the entire cohort and their genetic parents and we measured the transmission of variants to the next generation.

## Results

### Generating and sequencing CRISPR-Cas9-edited F0 and F1 individuals

We focused on three different genes (*anln, kmt2d*, and *smchd1*) for which (a) we have substantial experience in this model organism and (b) which give reproducible, quantitative defects in kidney morphogenesis (Hall et al., 2018), mandibular and neuronal development (Tsai et al., 2018), and craniofacial morphogenesis (Shaw et al., 2017). For each locus, we used sgRNAs that had the following three characteristics. First, for each of the three genes, we selected an sgRNA with demonstrated high efficiency (100%) and an sgRNA with low efficiency (∼30%), as determined by heteroduplex analysis and Sanger sequencing of cloned PCR products (Hall et al., 2018; Shaw et al., 2017; Tsai et al., 2018) (Suppl. Figure 1). Second, we mandated that all sgRNAs have a high specificity score (MIT specificity score 79-99 for each sgRNA; Suppl. Table S1). Finally, we required that each sgRNA was predicted to generate off-target effects with low cutting frequency determination (CFD) scores (mean = 0.17, range = 0-0.73; Suppl. Figure 2).

Next, we co-injected each sgRNA and Cas9 protein into wild-type zebrafish embryos from the same clutch at the 1-cell stage. For each sgRNA, we harvested DNA from six edited individuals to serve as technical replicates. In addition, we collected DNA from two individuals for each of the following conditions: uninjected, sgRNA alone, or Cas9 alone (Figure 1A). Finally, to assess the potential transmission of *de novo* variants to the next generation, we raised the F0 cohort for the *smchd1* high efficiency sgRNA and intercrossed adults to obtain the F1 generation. In total, we performed whole exome sequencing (WES) on two parents, 52 F0 individuals and 16 F1 individuals (Figure 1A). WES resulted in 76x average target coverage in F0 samples and 115x average target coverage in F1 individuals (Figure 1B, C). The F0 sequencing data covered 83% of the exome at ≥30x and 65% at ≥50x. The F1 sequencing data covered 88% of the exome at ≥30x and 78% of the exome at ≥50x.

**Figure 1.**
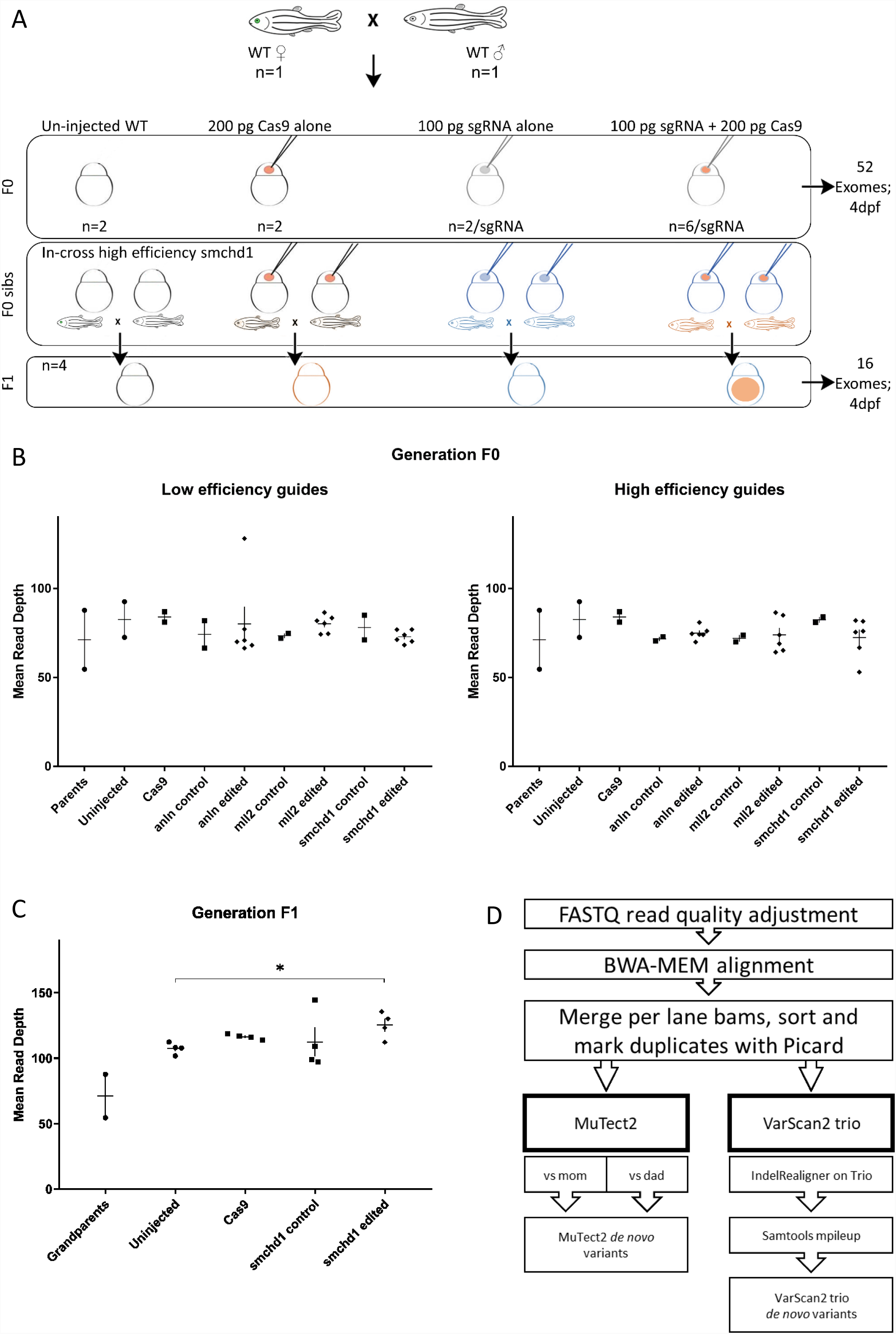
Whole exome sequencing in two generations of CRISPR-Cas9 edited zebrafish. (A) The experimental design generates a single clutch of embryos from a founder pair of parents from the ZDR laboratory strain of wild-type zebrafish. A total of 52 embryos were selected for DNA extraction and sequencing at 4 dpf in the F0 generation (2 uninjected, 2 Cas9 injected, 2 sgRNA injected across 6 different sgRNAs targeting 3 genes for a total of 12 embryos, and 6 CRISPR-Cas9 embryos per sgRNA guide for a total of 36 edited individuals). Additional embryos for each condition were injected concurrently, but raised to adulthood. The *smchd1* high efficiency guide F0 in-cross generated F1 progeny for further sequencing (4 uninjected, 4 Cas9 injected, 4 sgRNA injected, and 4 CRISPR-Cas9 injected embryos were selected for a total of 16 F1 exomes). (B) The first round of exome sequencing (F0 and parents) generated a consistent read depth averaging 76x coverage. (C) The second round of exome sequencing (F1) generated a consistently higher read depth averaging 115x coverage. The *smchd1* edited individuals are also sequenced to a higher depth than the uninjected controls (p<.05). (D) After sequencing quality control and alignment, variant calling was performed with both somatic and germline callers to identify candidate *de novo* mutations.

### *De novo* mutation counts are not inflated in F1 exomes

Low-level mosaicism remains challenging to detect in WES data and it is prone to high false-positive and false-negative rates (Sandmann et al., 2017). For this reason, we first focused on transmitted events. If CRISPR-Cas9 editing does induce off-target *de novo* mutations, we should observe an increase above baseline in the number of heterozygous variants fixed in the CRISPR-edited F1 generation that were absent from the grandparents.

Given the estimated 0.01% gene level baseline mutation rate in zebrafish(Mullins et al., 1994), we expect approximately 2-3 exonic changes per generation. To measure the observed rates, we applied a trio sequencing workflow aligned with GATK best practices and we called both single nucleotide variants and indels with two established variant callers: VarScan2 or Mutect2 (Figure 1D). Starting with all calls, we performed multiple data filtering steps. First, we removed variants present in either of the grandparental exomes. Second, since a small number of variants might have appeared *de novo* because of missing data from either grandparent, we also excluded alleles reported in the zebrafish ensembl dbSNP database. Third, we removed variants from the on-target genome locations (Suppl. Figure 3). Together, these three filters removed 79% of the MuTect2 and 99% of the VarScan2 calls. As an additional data filtering step, we removed repetitive elements, regions of potential segmental duplication in zebrafish, and indel variants containing homo-, dinucleotide, and trinucleotide repeats. This step improved the transition-transversion ratio from 0.91 to 1.09 which approaches a previously reported ratio of 1.2 for zebrafish (Stickney et al., 2002) (Suppl. Figure 4). Finally, we removed cross-noise variants found in two or more samples that likely represent systematic technical error or uncalled low-level mosaics from the grandparents.

Using this dataset (Suppl. Table S2), we then applied a filter for allele frequency (AF) above 0.3 to capture the fixed heterozygous variants and we compared the variant count differences between F1 embryos derived from edited and unedited F0 adults. VarScan2 reports candidate variant counts closer to the expected natural accumulation of *de novo* mutations in F1 than MuTect2 (average 20 vs 66, respectively; Suppl. Table S3). We calculated the critical p-value threshold Bonferroni correction for three groups (p<0.012), and neither calling method reports a significant difference between progeny of edited and control adults (p>0.11; Wilcox rank test, Suppl. Table S4).

Next, we focused on the VarScan2 results. Based on the >5-fold inflation of observed versus expected variant calls across the cohort (mean of 20 vs 2-3, respectively; Figure 2A) we hypothesized that these agnostically filtered calls still included false positives. Therefore, we reviewed the variant calls in the Integrative Genomics Viewer (IGV). We found two sources of false positives. First, a subset of read alignments filled into small deletions observed in the grandparents rather than extend a gap (83% of calls). Second, local realignments involving small deletions misalign in the progeny, even though an alternative placement of the deletion results in a grandparental genotype (10% of calls).

**Figure 2.**
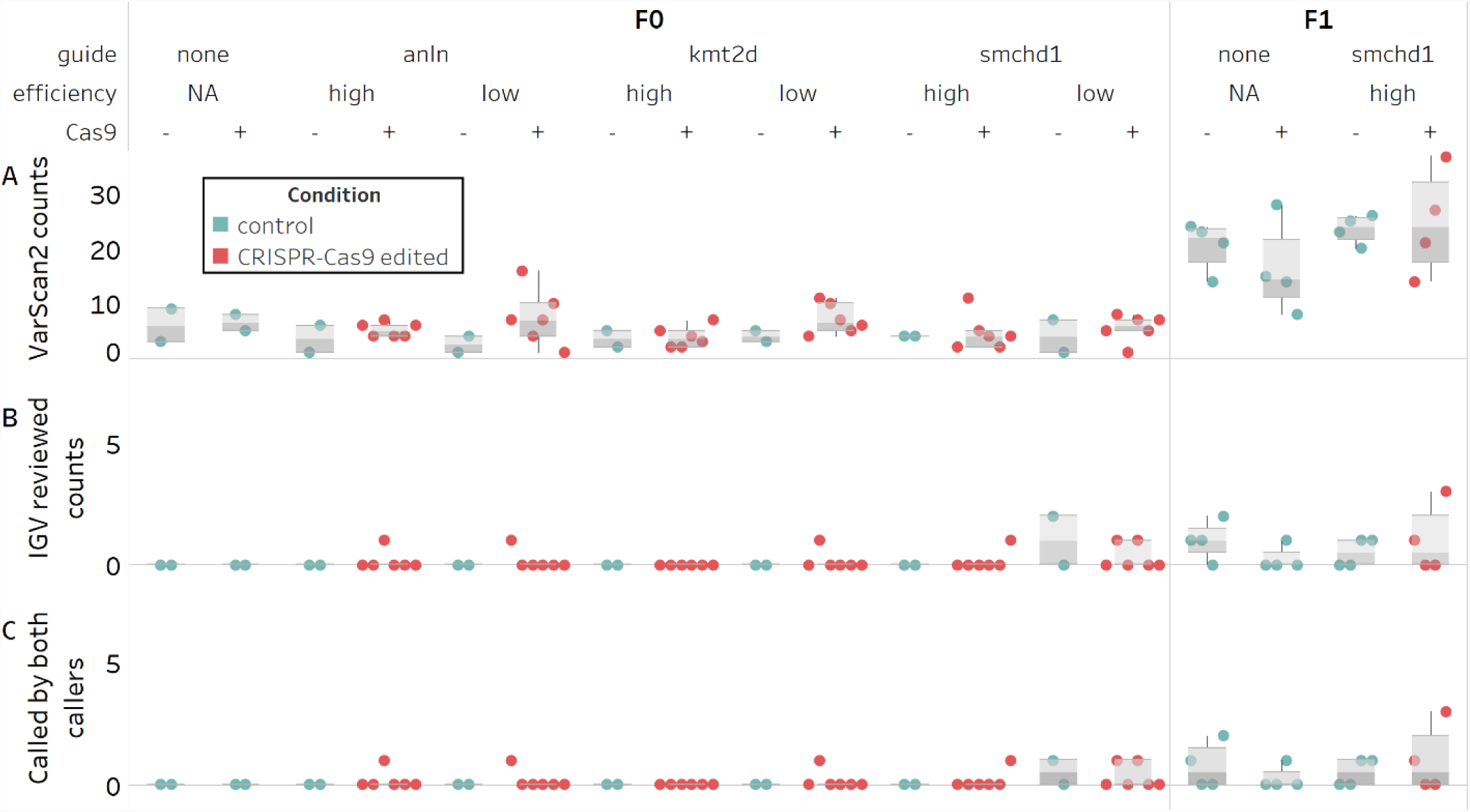
Counts of candidate *de novo* mutations in control and edited individual zebrafish embryos. Variants persisting after filtering and with an allele frequency ≥0.3 are not significantly different between control and CRISPR-Cas9 edited groups (N=68). (A) Predicted counts by VarScan2. (B) Unambiguous heterozygous variants determined by visual inspection of VarScan2 calls in IGV (C) Subset of predicted variants detected by both variants callers.

Of the remaining calls, half were deemed unlikely to be *bona fide* variants for other reasons. These included complex regions with many error prone reads; abundance of mis-mapped read pairs; and remaining low level mosaicism in grandparents. The other half were unambiguous *de novo* heterozygous variants (Figure 2B). Notably, most of the unambiguous variants were also called by MuTect2 (10 of 11; Figure 2C). For this population of alleles, we observed no difference between control and edited groups called by both callers (Suppl. Table S5). Crucially, we confirmed all of the variants detected by both callers in F1 animals derived from CRISPR/Cas edited individuals by Sanger sequencing. Taken together, we found that, regardless of whether we consider agnostic or manually reviewed variant numbers, there is no predilection toward inflated variant counts in F1 offspring derived from edited versus control groups. Further, the observed number of *de novo* variants in F1s does not exceed the expected rate of 2-3 per exome, per generation.

### *De novo* mutation counts are not inflated across the multigenerational cohort

We then returned to the F0 cohort to investigate whether variant burden outside of the targeted locus differed among individuals injected with sgRNA in the presence or absence of Cas9. Importantly, the expected allelic series of variants are reported robustly at the on-target locations of the sgRNAs against two of the target genes, *anln* on chromosome 19 and *kmt2d* on chromosome 23 (Supplementary Figure 3A) (Hall et al., 2018; Tsai et al., 2018). No on-target variants are observed for the *smchd1* locus because our exome capture did not include baits for this locus in the Zv9 assembly of the zebrafish genome. However, we demonstrated experimentally the on-target CRISPR-editing capability of the two *smchd1* sgRNAs and the transmission of on-target variants produced by the high-efficiency sgRNA to the F1 generation via Sanger sequencing (Supplementary Figure 3B), as described (Shaw et al., 2017).

We first considered the agnostic off-target VarScan2 variants called in the mosaic F0 generation (Suppl. Table S6). Initially, we applied the same arbitrary 0.3 AF threshold that we used with the F1 calls, reasoning that editing occurs at the one-to-two cell stage and would likely manifest as an off-target inflation at high allele frequencies. We determined the Bonferroni correction threshold for four groups (p<0.012), and again, we did not observe a significant inflation in *de novo* variant counts between control and F0 edited groups, in either the algorithmically predicted counts or the manually reviewed counts (p>0.15; Wilcox rank test; Figure 2A, B; Table S7). We then repeated the analysis on the agnostic MuTect2 call set, and consistent with the filtered VarScan2 data, we did not observe an inflation in *de novo* mutation counts between control and edited groups (p>0.04; Suppl. Table S7). Finally, because a 0.3 AF may fail to detect inefficient targeting events or lower mosaicism levels, we tested lower cutoff frequencies. At either an arbitrary 0.1 AF threshold, or without applying a threshold, we still observe no significant differences (p>0.08; Suppl. Table S7).

For the VarScan2 dataset generated from F0 exomes, the variant count exceeded the expected 2-3 *de novo* changes per exome in at least one individual in half of the edited conditions (Figure 2A). To exclude the possibility that these could be false positive calls, similar to what we observed in the F1 cohort, we inspected all variants exceeding the 0.3 AF cutoff using IGV. We found that this dataset also was subject to similar technical artifacts as observed for F1s; exclusion of these variants brought the *de novo* mutation call number within the expected range (Figure 2B). Using the same Bonferroni correction for four groups (p<0.012), we were unable to detect a difference between control versus edited groups (p>0.38; Suppl. Table S8). Since we had observed that variants detected by both callers represented an unbiased way to assess high confidence calls in F1, we also asked whether we could detect a difference in variant counts in this subset of calls in F0 (7 of 8 unambiguous calls; Figure 2C). Again, we observed no significant differences between controls and edited groups (p>0.78; Suppl. Table S9).

### *De novo* mutations are not observed at predicted off-target sites

To examine the potential incidence of off-target mutations more sensitively, we removed the filters on the variant calls and searched predicted off target sites across our multigenerational cohort using three algorithms: the MIT CRISPR design site, the CRISPR-direct engine, and CAS-OFFinder, for any variants occurring within 100 bp flanking a predicted off-target site. Consistent with previous reports (Hruscha et al., 2013; Varshney et al., 2015), we found no support for single nucleotide variants or small indels occurring at predicted off-target locations in the F1 generation, and sporadic low allele frequency calls near predicted off-target regions in F0s. The number of reported variants in the F0 samples are not significantly different than expected by chance (p>0.08; Supplementary Table S10).

We reviewed the 15 reported variant calls near predicted off-target sites in F0s, and found that none are supported by both variant callers (Supplementary Table S11). Seven are also reported in siblings subjected to editing with alternative guides or control conditions, making them unlikely to be induced by Cas9-mediated genome editing. Another four were not supported by reads on both strands. Of the four remaining variants, one was only reported in a control condition, making it unlikely to be a result of editing. The other three occur at a 5% alternate allele frequency, near the limit of detection for the variant callers, increasing the likelihood that they may be artifacts. We do note that one variant has features consistent with an expected off-target cut. This is a small deletion reported directly at a predicted off-target cut site detected by two prediction engines (Supplementary Table S10). Notably, this small deletion occurs in an exonic region, has a high CFD risk score (CFD score = 0.52), and is observed at the predicted locus in a few reads from the VarScan2 call set as well, even though it is not called by that algorithm. Together, our analysis of reported variants near predicted off-target sites detects one potential off-target variant at low allele frequency in a single individual and does not demonstrate an inflated or transmissible mutation burden conjoint with expected on-target deletions.

## Discussion

Trio sequencing designs enable off-target analyses to distinguish gene editing effects from natural and inherited genetic variation. In our study, the bulk of variant calls in zebrafish exomes are filtered out due to their existence in the parental strain. Our ability to recover transmissible on-target deletions and Sanger-validated *de novo* mutations outside of predicted off-target regions and in quantities indistinguishable from natural variation suggests that off-target CRISPR events occur infrequently.

Our results are consistent with previous results in zebrafish demonstrating limited off-target activity at select predicted regions (Hruscha et al., 2013; Varshney et al., 2015) and with recent work in mice that found limited support for off-target effects genome-wide (Iyer et al., 2018). However, we are limited to detecting potential off-target variation within the exon-capture space of the genome. We did not assess large structural variants or long deletions at the on-target site. In addition, we occasionally observed trends toward variant inflation in the predicted variant call sets that were related to sequencing depth and did not survive visual inspection. This observation suggests that even with trio designs and other precautionary measures, care should be exercised in interpreting variant predictions agnostically and that sequencing even more individuals per condition may be required to expose subtle differences in off-target effects.

In response to initial reports that CRISPR-Cas9 edited mammalian cells harbored off-target variants (Fu et al., 2013; Zhang et al., 2015), many iterative improvements in technology and experimental design have outlined conditions for achieving CRISPR-Cas9 gene editing while limiting off-target events. Our experimental and sgRNA design incorporated such advancements (high on-target MIT ranking, low off-target CFD scores, high cutting efficiency, and short Cas9 exposure), minimizing the chance of inducing off-target events. However, unexpected nuances of the CRISPR-Cas9 editing system continue to emerge. Varied biological responses to CRISPR-Cas9, such as DNA damage repair (Haapaniemi et al., 2018), enzymatic immunity (Crudele and Chamberlain, 2018), and alternative templating (Ma et al., 2017) exemplify our still nascent understanding of DNA and RNA editing. Furthermore, natural human genetic variation has been shown to influence both the efficaciousness of on-target editing and the frequency of off-target editing (Lessard et al., 2017). Under these circumstances, use of emergent computational, laboratory, and animal modeling tools and unbiased genome-wide off-target assessments will facilitate the foundational knowledge required to reduce unnecessary risk in practice.

## Methods

### CRISPR-Cas9 gene editing in zebrafish embryos

We used CHOPCHOP (Labun et al., 2016) to identify sgRNAs targeting a sequence within the coding regions of the target genes and sgRNAs were *in vitro* transcribed using the GeneArt precision gRNA synthesis kit (Thermo Fisher, Waltham, MA) according to the manufacturer’s instructions. See Supplemental Figure S1, Table S1, and references (Hall et al., 2018; Shaw et al., 2017; Tsai et al., 2018) for details on targeting sequences/locations and sgRNA efficiency. Zebrafish embryos from a single clutch from a natural mating of a ZDR background founder pair were either uninjected or injected into the cell at the 1-cell stage with a 1 nl cocktail of 100 pg/nl sgRNA, 200 pg/nl Cas9 protein (PNA Bio, Newbury Park, CA), or a combination of both reagents. We extracted genomic DNA (gDNA) from tail clips of parental zebrafish or whole zebrafish embryos at 4 dpf.

### Sample Selection for Sequencing

The ZDR strain in our laboratory gives consistently robust clutch sizes of ∼100 embryos. To preserve enough individuals to generate an F1 generation, we anticipated that we would have approximately 50 individuals available for exome sequencing. Using the CFD score cut-off of 0.2 as a threshold for the likelihood of inducing transmissible off-target mutations, we expected that we would need at least 5-6 embryos per condition to observe one of these events. Thus, we selected six independent embryos per gRNA plus Cas9 condition for comparison with controls while maintaining the experiment within a single clutch to control for inherited variation.

### Heteroduplex Editing Efficiency by PAGE

For each sgRNA plus Cas9 condition we PCR-amplified gDNA from 12 embryos per batch using site-specific primers and screened for heteroduplex formation as described (Zhu et al., 2014). Five samples with evidence of heteroduplex formation were gel purified alongside a control sample, ‘A’ overhangs were added to the PCR products, and the products were cloned into a TOPO4 vector (Thermo Fisher). We picked 12 colonies per embryo to estimate targeting efficiency by Sanger sequencing.

### Whole Exome Sequencing

We used the manufacturer protocol for the Agilent SureSelect Capture kit for non-human exomes with 200 ng gDNA per individual (75 Mb capture designed on the zv9 version of the zebrafish genome; Agilent SSXT Zebrafish All Exon kit; Agilent Technologies, Santa Clara, CA). Samples were multiplexed and run across two lanes of the Illumina HiSeq 4000 as paired-end 150 bp reads. Sequence data were demultiplexed and Fastq files were generated using Bcl2Fastq conversion software (Illumina, San Diego, CA).

### Variant Calling

Sequencing reads were processed using the TrimGalore toolkit (Krueger, 2017) which employs Cutadapt to trim low quality bases and Illumina sequencing adapters from the 3’ end of the reads. Only reads that were 20 nt or longer after trimming were kept for further analysis. Using the BWA (v. 0.7.15) MEM algorithm (Li, 2013), reads were mapped to the Zv9 version of the zebrafish genome. Picard tools (*Picard*, 2017) (v. 2.14.1) were used to remove PCR duplicates and to calculate sequencing metrics. The Genome Analysis Toolkit(McKenna et al., 2010) (GATK, v. 3.8-0) MuTect2 caller was used to call variants between each experimental condition and the adult male and adult female samples separately. Independently, aligned reads were locally realigned with the GATK IndelRealigner and then processed with Samtools mpileup(Li, 2011) for variant calling with VarScan2 trio (Koboldt et al., 2013). VarScan2 variant call sets were generated with the minimum coverage specified at 30x.

### Variant analysis

We used BEDOPS (Neph et al., 2012) and Bedtools (Quinlan and Hall, 2010) intersect, window, and merge commands to exclude variants with support in either parent, variants reported to occur in wild-type zebrafish strains ensembl dbSNP version 79, variants in repeat regions or regions of predicted segmental duplication in the genome (Khaja et al., 2006), variants reported in both control individuals and CRISPR-edited individuals, and variants reported at the on-target locations for CRISPR-editing. The potential for variants to occur due to off-target CRISPR-mediated editing was assessed by comparing variant counts between groups with either a Wilcoxon rank test for two groups, or a Kruskal-Wallis rank test for more than two groups and assessing the p-value against a Bonferroni critical value to correct for multiple testing. In addition, variants from samples were compared with locations of predicted off-target regions (formatted into a .bed file) from three algorithms: CRISPOR (Concordet and Haeussler, 2018), the CRISPRdirect engine with 12-mer to 20-mer hits, or Cas9-OFFinder allowing 3-mismatches and 1-bulge in either DNA or RNA. Hypergeometric p-values calculated with the Rothstein lab hypergeometric calculator, use the capture space (74691693 bp) as the population size, and a reasonable high vs low sequencing error rate for our Illumina platform (.24% vs .1%) (Pfeiffer et al., 2018) to calculate the expected number of population variants called by chance at a position covered at the F0 average read depth (4 or more errant reads at the position; AF >.05).

## Acknowledgements

We are grateful to I-Chun Tsai, Maria Kousi, Zachary Kupchinsky and Igor Pediaditakis for technical assistance. This work was supported by a fellowship from U.S. National Institutes of Health Grant 5T32HG008955-02 (M.M.). We thank Nicolas Devos (Duke Sequencing and Genomic Technologies Shared Resource) and David Corcoran (Duke Genomic Analysis and Bioinformatics Shared Resource) for sequencing and informatics support, respectively. Some analyses were carried out using resources from the Duke Compute Cluster. N.K. is a Distinguished Jean and George Brumley Professor.

---

**Figure S1.**
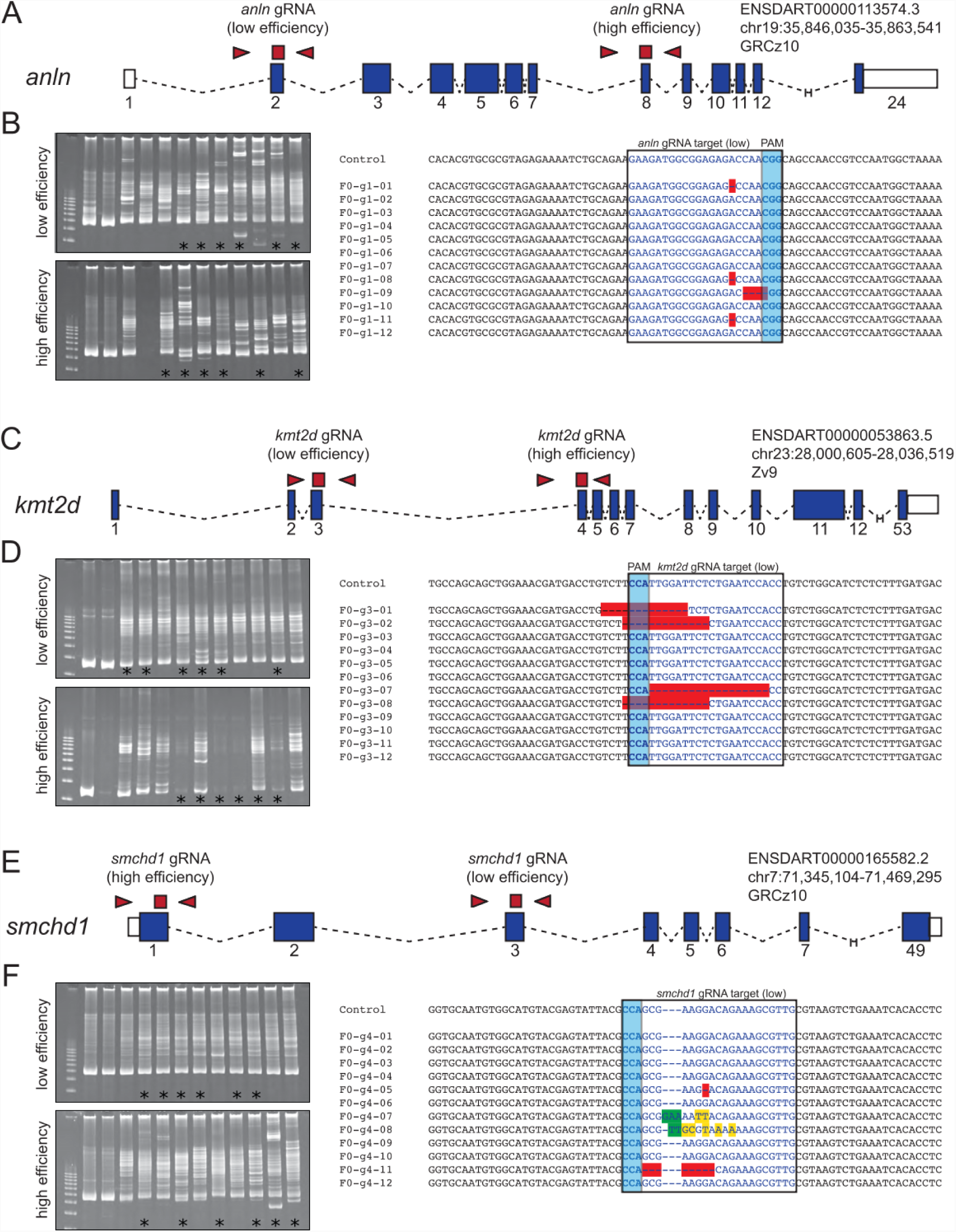
Confirmation of CRISPR editing efficiencies. Efficiency data for the high efficiency guides have been published previously. (A, C, E) Schematic of the *D. rerio* locus, sgRNA targeted regions (red squares) and primers used to determine sgRNA efficiency (red triangles) for each gene of interest. (B, D, F) Heteroduplex analysis (left) and Sanger sequencing of 12 clones amplified from a single representative embryo injected with the low efficiency sgRNA plus Cas9 for each target gene (right). Efficiency was estimated by taking the average number of targeted clones across five embryos per sgRNA. * denotes samples from the heteroduplex analysis choosen for sequencing; PAM, protospacer adjacent motif.

**Figure S2.**
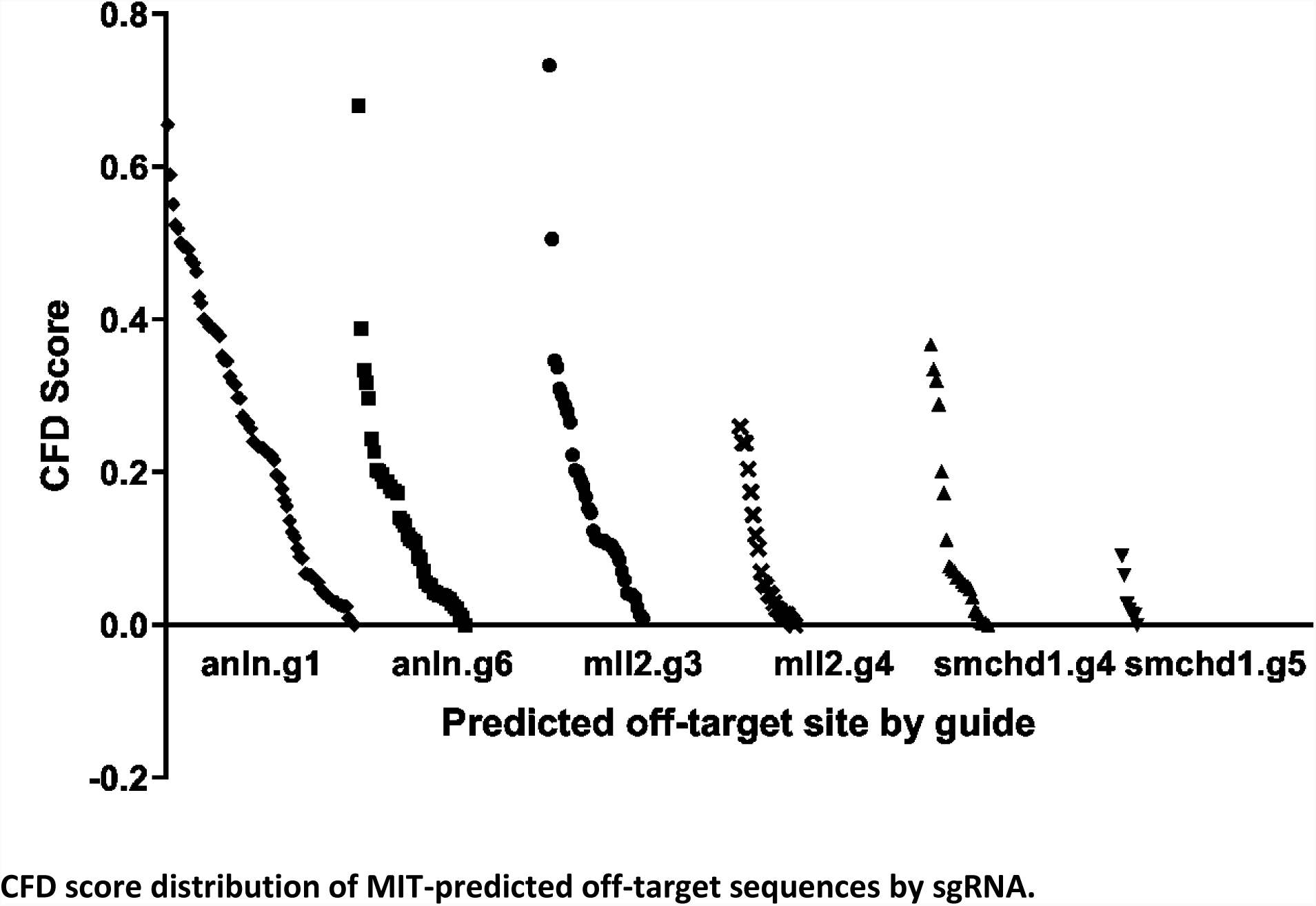
CFD score distribution of MIT-predicted off-target sequences by sgRNA.

**Figure S3.**
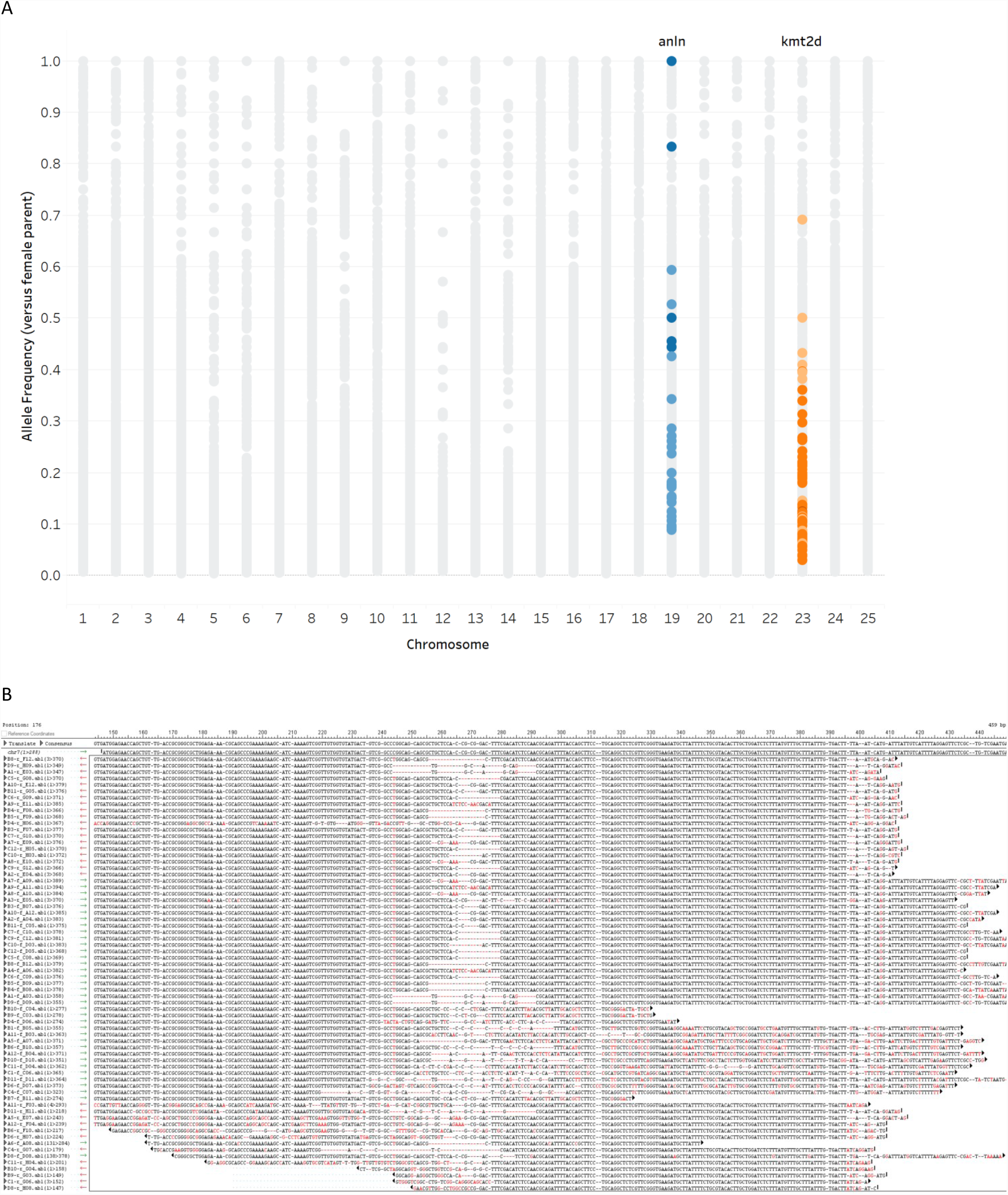
On-target germline CRISPR-Cas9 editing transmitted to F1. (A) Grey circles represent all variants calls. On-target allelic series at the *anln* locus (blue) and *kmt2d* locus (orange). (B) Sequencing at the on-target *smchd1* locus in F1s originating from F0s injected with high-efficiency sgRNA plus Cas9.

**Figure S4.**
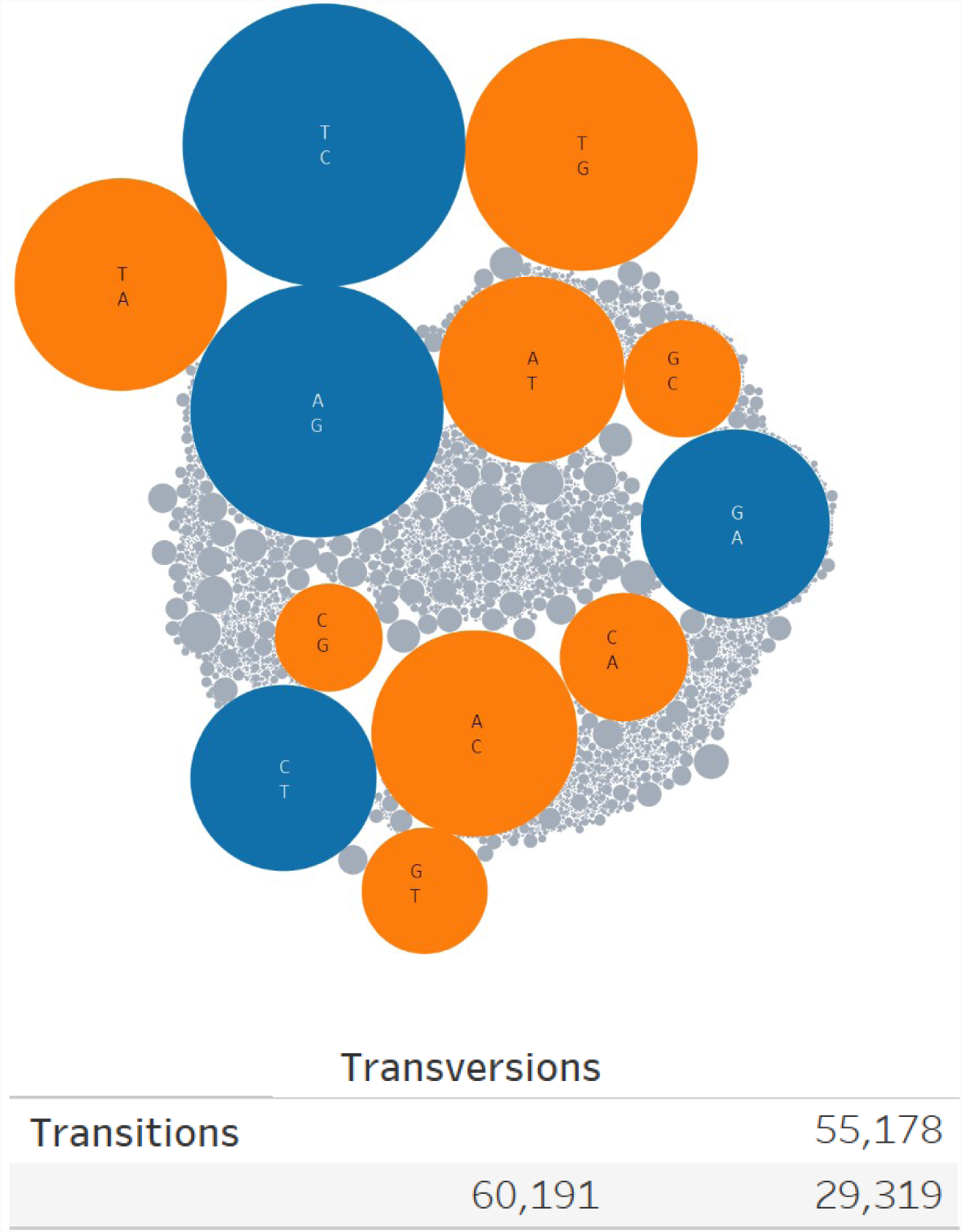
Transition-Transversion ratio in F1 exomes compared to grandparental exomes. Sizes of circles represent the number of observations for each variant class: transitions (blue), transversions (orange), indels (grey). After filtering, the transition-transversion ratio is 1.09.

**Table S1:**
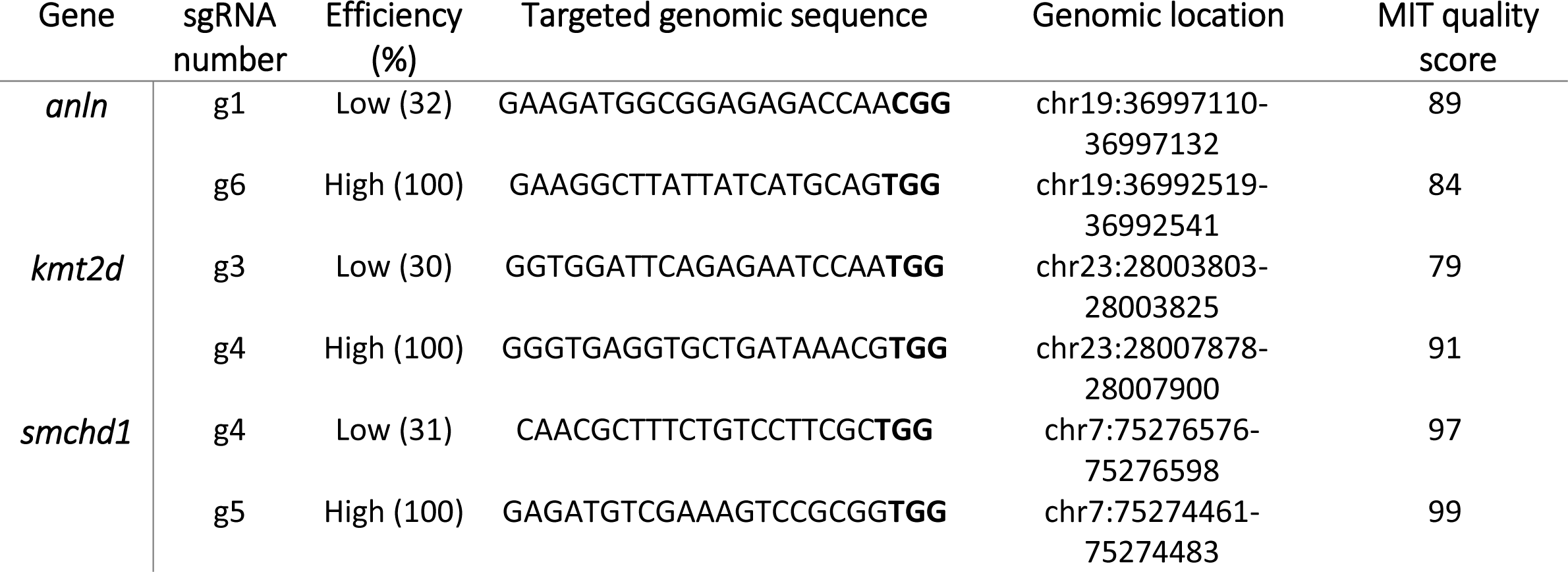
sgRNA gene targets, efficiencies, and quality scores.

**Table S2:** Variant counts during filtering in F1 individuals.

| Sample | MuTect2 |  |  | VarScan2 |  |  |
| --- | --- | --- | --- | --- | --- | --- |
| | originally reported* | exclude grand-parent & dbSNP | passing filters + AF $\geq 0.3$ | originally reported | exclude grand-parent & dbSNP | passing filters + AF $\geq 0.3$ |
| smchd1-S1 | 107784 | 16467 | 93 | 721082 | 4836 | 23 |
| smchd1-S2 | 91320.5 | 14343 | 65 | 720522 | 5444 | 14 |
| smchd1-S3 | 93910 | 14820 | 75 | 722006 | 5175 | 23 |
| smchd1-S4 | 98067 | 16162 | 77 | 721454 | 4983 | 19 |
| smchd1-Cas9-S5 | 99133.5 | 15695 | 70 | 722267 | 5152 | 28 |
| smchd1-Cas9-S6 | 98879 | 18379 | 47 | 722224 | 5048 | 13 |
| smchd1-Cas9-S7 | 90394.5 | 16366 | 79 | 722826 | 5074 | 15 |
| smchd1-Cas9-S8 | 102871.5 | 15899 | 48 | 722018 | 4847 | 8 |
| smchd1-g5-S9 | 111667.5 | 22892 | 82 | 721814 | 5790 | 25 |
| smchd1-g5-S10 | 93602.5 | 13941 | 51 | 718963 | 5279 | 19 |
| smchd1-g5-S11 | 102256.5 | 12190 | 86 | 718274 | 4909 | 23 |
| smchd1-g5-S12 | 100041 | 14611 | 73 | 718720 | 5157 | 25 |
| smchd1-g5-Cas9-S13 | 96334.5 | 18489 | 52 | 721700 | 5327 | 21 |
| smchd1-g5-Cas9-S14 | 91999 | 18250 | 63 | 720369 | 5059 | 27 |
| smchd1-g5-Cas9-S15 | 98719 | 18893 | 46 | 720855 | 5252 | 13 |
| smchd1-g5-Cas9-S16 | 97144.5 | 17803 | 70 | 720601 | 5114 | 34 |

\*Reported count is an average of the counts called against the female grandparent and the counts called against the male grandparent.

**Table S3:** Average count of candidate *de novo* mutations by generation and calling method.

|  |  |  |  | MuTect2 |  | VarScan2 |  |
| --- | --- | --- | --- | --- | --- | --- | --- |
| sgRNA | Efficiency | Cas9 | Condition | F0 | F1 | F0 | F1 |
| none |  | absent | control | 41 | 76 | 6 | 19 |
|  |  | present | control | 24 | 60 | 6 | 16 |
| smchd1 | high | absent | control | 21 | 72 | 4 | 23 |
|  |  | present | edited | 26 | 55 | 4 | 23 |
|  | low | absent | control | 27 |  | 3 |  |
|  |  | present | edited | 22 |  | 5 |  |
| anln | high | absent | control | 19 |  | 3 |  |
|  |  | present | edited | 29 |  | 5 |  |
|  | low | absent | control | 13 |  | 2 |  |
|  |  | present | edited | 33 |  | 7 |  |
| kmt2d | high | absent | control | 23 |  | 3 |  |
|  |  | present | edited | 27 |  | 3 |  |
|  | low | absent | control | 22 |  | 4 |  |
|  |  | present | edited | 39 |  | 7 |  |
| Average |  |  |  | 26 | 66 | 4 | 20 |

**Table S4:** Wilcox comparison p-values in F1.

| Sample | MuTect2 | Varscan2 |
| --- | --- | --- |
| Control vs edited | 0.11 | 0.47 |
| Cas9 present vs absent | 0.03 | 0.71 |
| Guide present vs absent | 0.64 | 0.15 |

**Table S5:** Wilcox comparison p-values on variants called by both callers in F1.

| Sample | AF ≥ 0.3 |
| --- | --- |
| Control vs edited | 0.78 |
| Cas9 present vs absent | 0.57 |
| Guide present vs absent |  |
| High vs none | 1.0 |

**Table S6:** Variant counts during filtering in F0 individuals.

| Sample | MuTect2 |  |  | ≥30x VarScan2 |  |  |
| --- | --- | --- | --- | --- | --- | --- |
|  | originally reported* | exclude parent & dbSNP | passing filters + AF ≥ 0.3 | originally reported | exclude parent & dbSNP | passing filters + AF ≥ 0.3 |
| uninjected-3 | 98730 | 18231 | 19 | 714178 | 4281 | 3 |
| uninjected-5 | 153632.5 | 38063 | 63 | 724678 | 6727 | 9 |
| Cas9-4 | 133284 | 25688 | 25 | 719479 | 4405 | 8 |
| Cas9-5 | 133735 | 25761 | 24 | 713112 | 4663 | 5 |
| anIn-g1-1 | 117437.5 | 16151 | 12 | 706727 | 4899 | 1 |
| anIn-g1-3 | 140097 | 28430 | 15 | 720890 | 5970 | 4 |
| anIn-g1-Cas9-10 | 125951.5 | 19246 | 26 | 706488 | 4593 | 1 |
| anIn-g1-Cas9-4 | 109913 | 21248 | 23 | 712796 | 4419 | 4 |
| anIn-g1-Cas9-5 | 125363.5 | 27518 | 23 | 717865 | 5235 | 7 |
| anIn-g1-Cas9-6 | 152948.5 | 43391 | 84 | 726997 | 7228 | 16 |
| anIn-g1-Cas9-7 | 123259.5 | 19800 | 25 | 711674 | 4780 | 10 |
| anIn-g1-Cas9-9 | 113957.5 | 16139 | 19 | 704081 | 4497 | 6 |
| anIn-g6-3 | 122569.5 | 26063 | 17 | 712519 | 4386 | 1 |
| anIn-g6-4 | 131988.5 | 22096 | 21 | 710221 | 4236 | 6 |
| anIn-g6-Cas9-10 | 139893.5 | 30275 | 21 | 718040 | 5273 | 6 |
| anIn-g6-Cas9-3 | 116501 | 26021 | 36 | 713070 | 4956 | 6 |
| anIn-g6-Cas9-4 | 123799.5 | 26379 | 48 | 717711 | 5671 | 4 |
| anIn-g6-Cas9-5 | 132917.5 | 25629 | 28 | 715359 | 5025 | 4 |
| anIn-g6-Cas9-6 | 137644 | 29799 | 19 | 715692 | 4722 | 4 |
| anIn-g6-Cas9-8 | 120272.5 | 24997 | 25 | 719710 | 4659 | 6 |
| kmt2d-g3-4 | 116601.5 | 20554 | 22 | 722869 | 5663 | 5 |
| kmt2d-g3-5 | 139178 | 28685 | 23 | 714383 | 5739 | 3 |
| kmt2d-g3-Cas9-1 | 118623 | 24801 | 60 | 720884 | 4646 | 10 |
| kmt2d-g3-Cas9-2 | 116798 | 20458 | 46 | 715781 | 5446 | 7 |
| kmt2d-g3-Cas9-4 | 133515.5 | 34500 | 45 | 722333 | 5807 | 10 |
| kmt2d-g3-Cas9-5 | 111948.5 | 25153 | 32 | 715519 | 4491 | 5 |
| kmt2d-g3-Cas9-6 | 119867.5 | 24248 | 28 | 719554 | 5316 | 4 |
| kmt2d-g3-Cas9-9 | 133272 | 30346 | 23 | 720748 | 5624 | 6 |
| kmt2d-g4-1 | 123010.5 | 31250 | 26 | 716594 | 5504 | 5 |
| kmt2d-g4-5 | 124993.5 | 26831 | 21 | 713325 | 5172 | 2 |
| kmt2d-g4-Cas9-4 | 119952.5 | 24033 | 28 | 707322 | 4837 | 2 |
| kmt2d-g4-Cas9-5 | 119101.5 | 22956 | 20 | 705218 | 4510 | 4 |
| kmt2d-g4-Cas9-6 | 122240 | 26406 | 31 | 711825 | 5501 | 2 |
| kmt2d-g4-Cas9-7 | 119373 | 33534 | 31 | 717861 | 5671 | 3 |
| kmt2d-g4-Cas9-8 | 137530 | 33943 | 23 | 722359 | 5524 | 5 |
| kmt2d-g4-Cas9-9 | 134129 | 37854 | 32 | 721268 | 5812 | 7 |
| smchd1-g4-1 | 128430.5 | 26487 | 32 | 721553 | 5109 | 6 |
| smchd1-g4-2 | 122734.5 | 32737 | 22 | 714473 | 5452 | 1 |
| smchd1-g4-Cas9-2 | 112519.5 | 21120 | 18 | 712893 | 4548 | 1 |
| smchd1-g4-Cas9-3 | 133350.5 | 28182 | 31 | 715977 | 4646 | 6 |
| smchd1-g4-Cas9-4 | 129172.5 | 26522 | 17 | 709164 | 4852 | 7 |
| smchd1-g4-Cas9-5 | 120364 | 26252 | 24 | 716579 | 4391 | 5 |
| smchd1-g4-Cas9-7 | 125670.5 | 24490 | 28 | 718073 | 4766 | 5 |
| smchd1-g4-Cas9-8 | 124193.5 | 26396 | 17 | 713050 | 4667 | 7 |
| smchd1-g5-2 | 139507 | 34868 | 16 | 719051 | 5491 | 4 |
| smchd1-g5-3 | 133208.5 | 32225 | 27 | 721170 | 5669 | 4 |
| smchd1-g5-Cas9-10 | 125004 | 27506 | 13 | 707974 | 4411 | 3 |
| smchd1-g5-Cas9-2 | 128403 | 29684 | 33 | 716875 | 4833 | 5 |
| smchd1-g5-Cas9-4 | 127675.5 | 27174 | 24 | 717089 | 5516 | 4 |
| smchd1-g5-Cas9-6 | 126063 | 31473 | 42 | 721368 | 5204 | 11 |
| smchd1-g5-Cas9-8 | 95182.5 | 15438 | 14 | 668596 | 3244 | 2 |
| smchd1-g5-Cas9-9 | 107702 | 24169 | 33 | 721002 | 6121 | 2 |

\*Reported count is an average of the counts called against the female parent and the counts called against the male parent.

**Table S7:** Wilcox comparison p-values on predicted variants in F0.

| Sample | MuTect2 |  |  | VarScan2 |  |  |
| --- | --- | --- | --- | --- | --- | --- |
|  | AF ≥ 0.3 | AF ≥ 0.1 | All calls | AF ≥ 0.3 | AF ≥ 0.1 | All calls |
| Control vs edited | 0.04 | 0.08 | 0.89 | 0.15 | 0.48 | 0.82 |
| Cas9 present vs absent | 0.04 | 0.15 | 0.74 | 0.05 | 0.34 | 0.89 |
| Guide present vs absent |  |  |  |  |  |  |
| High vs none | 0.90 | 0.31 | 0.58 | 0.16 | 0.26 | 0.41 |
| Low vs none | 0.69 | 0.86 | 0.87 | 0.69 | 0.49 | 0.58 |

**Table S8:**
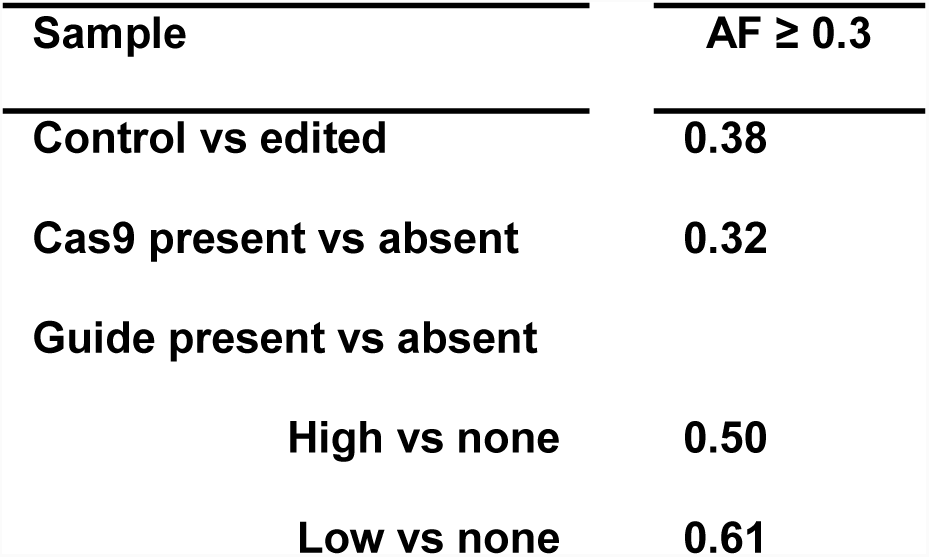
Wilcox comparison p-values on reviewed variants in F0 (VarScan2 only)

**Table S9:** Wilcox comparison p-values on variants called by both callers in F0.

| Sample | AF ≥ 0.3 | AF ≥ 0.1 | No filter |
| --- | --- | --- | --- |
| Control vs edited | 0.78 | 0.49 | 0.97 |
| Cas9 present vs absent | 0.96 | 0.77 | 0.51 |
| Guide present vs absent |  |  |  |
| High vs none | 1.0 | 1.0 | 0.25 |
| Low vs none | 0.89 | 0.86 | 0.44 |

**Table S10:** Expected off-target observations in region by chance, hypergeometric p-value.

| **Guide** | **Sample size**<br>**(# off-target sites**<br>**x 200bp)** | **Max.**<br>**observed**<br>**variants in**<br>**individual** | **p-value (under-**<br>**represented) vs**<br>**population max.**<br>**variant rate** | **p-value (under-**<br>**represented) vs**<br>**population min.**<br>**variant rate** |
| --- | --- | --- | --- | --- |
| **anln-g1** | 317200 | 2 | 1.07e-17 | .531 |
| **anln-g6** | 524400 | 1 | 5.63e-32 | .077 |
| **kmt2d-g3** | 269200 | 1 | 4.2e-16 | .363 |
| **kmt2d-g4** | 169200 | 1 | 5.5e-10 | .606 |
| **smchd1-g4** | 183400 | 0 | NA | NA |
| **smchd1-g5** | 151400 | 1 | .367 | .657 |

Hypergeometric p-values calculated with the Rothstein lab hypergeometric calculator, using the capture space (74691693 bp) as the population size, and a high or low estimate for the expected population variant counts (10975 vs 608, respectively).

**Table S11:** Variants at predicted off-target loci.

| Sample | Condition | Variant | Ref | Alt | Variant<br>Caller | Predicted<br>off-target | Prediction<br>algorithm | Allele<br>Frequency |
| --- | --- | --- | --- | --- | --- | --- | --- | --- |
| anln-g1-3 | Guide only | 24:28102298* | A | C | VarScan2 | 24:28102339-61 | Cas-OFFinder | 0.05 |
| anln-g1-Cas9-7 | Edited | 24:28102298* | A | C | VarScan2 | 24:28102339-61 | Cas-OFFinder | 0.05 |
| anln-g1-3 | Guide only | 2:44898792** | C | T | VarScan2 | 2:44898891-913 | MIT CRISPOR | 0.06 |
| anln-g1-Cas9-9 | Edited | 2:44898798** | G | T | VarScan2 | 2:44898891-913 | MIT CRISPOR | 0.06 |
| anln-g1-Cas9-6 | Edited | 7:65593684** | AC | A | VarScan2 | 7:65593592-602 | CRISPR-direct | 0.1 |
| anln-g6-Cas9-10 | Edited | 16:54853302-9** | Clustered event |  | MuTect2 | 16:54853401-11 | CRISPR-direct | <0.05 |
| kmt2d-g4-Cas9-7 | Edited | 5:62586647** | A | ACAC | VarScan2 | 5:62586546-56 | CRISPR-direct | 0.26 |
| anln-g6-Cas9-3 | Edited | 16:10603230† | C | A | VarScan2 | 16:10603172-82 | CRISPR-direct | 0.06 |
| kmt2d-g3-Cas9-1 | Edited | 22:20459578† | C | T | VarScan2 | 22:20459646-56 | CRISPR-direct | 0.08 |
| kmt2d-g3-Cas9-2 | Edited | 22:20459578† | C | T | VarScan2 | 22:20459646-56 | CRISPR-direct | 0.08 |
| kmt2d-g3-Cas9-4 | Edited | 22:20459578† | C | T | VarScan2 | 22:20459646-56 | CRISPR-direct | 0.1 |
| smchd1-g5-3 | Guide only | 2:50824997-9 | CTG | AAA | MuTect2 | 2:50824943-53 | CRISPR-direct | 0.05 |
| kmt2d-g4-Cas9-5 | Edited | 14:2529163 | A | G | VarScan2 | 14:2529206-28 | Cas-OFFinder | 0.05 |
| smchd1-g5-Cas9-2 | Edited | 25:21049069-77 | ACTAGCGTG | A | MuTect2 | 25:21048965-75 | CRISPR-direct | <0.05 |
| anln-g1-Cas9-6 | Edited | 14:11156689 | GAGACC | A | MuTect2 | 14:11156666-99 | MIT<br>CRISPOR,<br>Cas-OFFinder | <0.05 |

\*Variant at off-target site reported in both control and edited samples for this guide
\*\*Variant observed in siblings from other conditions
† Variant at off-target site not supported by reads on both strands

